# Rapid and quantitative detection of COVID-19 markers in micro-liter sized samples

**DOI:** 10.1101/2020.04.20.052233

**Authors:** Xiaotian Tan, Cory Lin, Jie Zhang, Maung Kyaw Khaing Oo, Xudong Fan

## Abstract

COVID-19 pandemic has caused tens of thousands of deaths and is now a severe threat to global health. Clinical practice has demonstrated that the SARS-CoV-2 S1 specific antibodies and viral antigens can be used as diagnostic and prognostic markers of COVID-19. However, the popular point-of-care biomarker detection technologies, such as the lateral-flow test strips, provide only yes/no information and have very limited sensitivities. Thus, it has a high false negative rate and cannot be used for the quantitative evaluation of patient’s immune response. Conventional ELISA (enzyme-linked immunosorbent assay), on the other hand, can provide quantitative, accurate, and sensitive results, but it involves complicated and expensive instruments and long assay time. In addition, samples need to be sent to centralized labs, which significantly increases the turn-around time. Here, we present a microfluidic ELISA technology for rapid (15-20 minutes), quantitative, sensitive detection of SARS-CoV-2 biomarkers using SARS-CoV-2 specific IgG and viral antigen – S protein in serum. We also characterized various humanized monoclonal IgG, and identified a candidate with a high binding affinity towards SARS-CoV-2 S1 protein that can serve as the calibration standard of anti-SARS-CoV-2 S1 IgG in serological analyses. Furthermore, we demonstrated that our microfluidic ELISA platform can be used for rapid affinity evaluation of monoclonal anti-S1 antibodies. The microfluidic ELISA device is highly portable and requires less than 10 μL of samples for each channel. Therefore, our technology will greatly facilitate rapid and quantitative analysis of COVID-19 patients and vaccine recipients at point-of-care.

## Introduction

The disease (COVID-19) related to novel coronavirus (SARS-CoV-2) has caused tens of thousands of deaths and is becoming a severe threat to global health. However, diagnosis and prognosis of COVID-19 are still facing serious challenges. The current “gold standard” method to diagnose SARS-CoV-2 infection is based on the detection of viral RNA (N gene) in nasal swab samples with RT-PCR. However, PCR-based diagnosis is prone to false negatives due to the uncertainties in sample collection and the molecular mechanism of the test and is unable to track the immune response to the viral infection^1^. Earlier SARS-CoV related research^2, 3^ and most recent studies on SARS-CoV-2^4, 5^ have shown that SARS-specific antibodies, such as IgG, IgM, and IgA, can be detected in serum as early as seven days after viral infection (3-5 days after the symptoms appear) and can last for several years after recovery^1^. Similar immune response should also occur after a successful vaccination (>7 days after vaccination)^2, 3, 6^. Therefore, virus-specific antibodies in serum and secretory mucus (*e.g*., saliva and sputum) can be used as diagnostic markers for viral infection and for the evaluation of patient’s adaptive immune responses (either from virus infection or from vaccination). As such, SARS-CoV-2-specific antibodies are currently listed among the diagnostic markers in the “COVID-19 Diagnosis and Treatment Plan (Provisional 7^th^ Edition)” published in China. In addition to SARS-CoV-2 specific antibodies, the viral antigen (such as the spike protein or S protein) in circulating blood can be used for the prognosis of COVID-19-related viremia^1, 7^. As one of the most commonly used targets for vaccine development, the serum concentration of the S protein is also a potential marker for early-stage vaccine responses, especially for sub-unit vaccines ^8^.

Unfortunately, the existing antibody detection methods are still far from adequate. The gold nanoparticle based lateral flow assay (*e.g*., paper-based test strips) is very popular in rapid detection of IgG/IgM antibodies (especially for point-of-care diagnostics)^9-14^. Although fast (5-20 minutes), it provides only yes/no information and has very limited sensitivities, making this method very easy to generate false positives/negative and impossible to track the patients’ immune response to infection, treatment, and vaccination. Conventional ELISA (enzyme-linked immunosorbent assay), on the other hand, can provide quantitative, accurate, and sensitive results, but it involves complicated and expensive instruments and long assay time (2-3 hours)^5, 15^. In addition, samples need to be sent to centralized labs, which significantly increases the turn-around time.

In this work, we present a microfluidic ELISA technology for rapid (15-20 minutes), quantitative, and sensitive detection of SARS-CoV-2 biomarkers using SARS-CoV-2 specific IgG and viral antigen – S protein, both of which are spiked in serum, as a model system. We also characterized various humanized monoclonal IgG and identified one with a high binding affinity towards SARS-CoV-2 S1 protein that can serve as the calibration standard of anti-SARS-CoV-2 S1 IgG in serological analyses. Furthermore, we demonstrated that our microfluidic ELISA platform can be used for rapid affinity evaluation of monoclonal anti-S1 antibodies. The microfluidic ELISA device is highly portable and requires only 10 μL of samples for each channel, which can be easily collected from a drop of fingertip blood. Therefore, our technology will greatly facilitate rapid and quantitative analysis of COVID-19 patients and vaccine recipients at point-of-care.

## Materials and methods

### Automated ELISA System

A detailed description of the automated ELISA system and corresponding capillary sensor arrays can be found in our previous publications^16, 17^. A photo of the capillary sensor array can be found in Figure 1(A). It is made of polystyrene using the injection molding method. The sensor array has 12 channels, each of which has an inner diameter of 0.8 mm and approximately 8 μL volume, and acts as an ELISA reactor.

**Figure 1.**
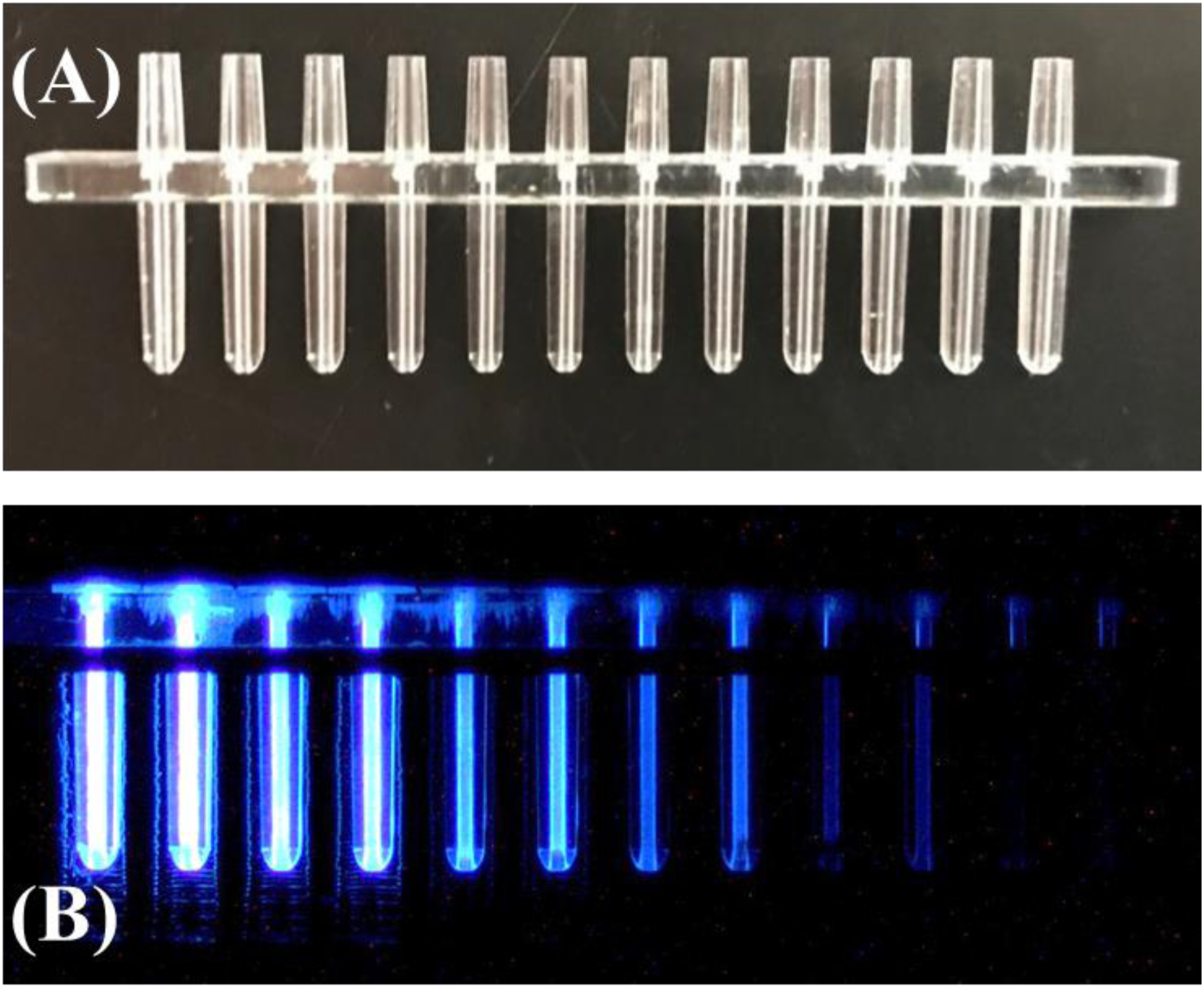
(A) Picture of a 12-channel disposable polystyrene cartridge. (B) Chemiluminescence signal from the cartridge for detection of various concentrations of analytes.

### Chemical Reagents

The chemiluminescent substrate (SuperSignal™ ELISA Femto Substrate, 37075), the UltraPure™ DNase/RNase-free distilled water (10977023), and the SuperBlock™ (PBS) blocking buffer (37515) were purchased from Thermo Fisher. The ELISA coating buffer (1× PBS, DY006), concentrated wash buffer (WA126), and concentrated reagent diluent (10% BSA in 10× PBS, DY995) were purchased from R&D Systems. The human serum was purchased from Millipore Sigma (H4522-20ML). Human-cell-expressed SARS-CoV-2 Spike S1-His recombinant protein was provided by Sino Biological (40591-V08H). The human-cell-expressed SARS-CoV Spike S1-His recombinant protein was purchased from Creative Biolabs (VAng-Wyb7337). The recombinant CR3022 therapeutic antibody was purchased from Creative Biolabs (MRO-1214LC). The humanized chimeric antibodies D001, D003, and D006 were developed and provided by Sino Biological (Catalog number: 40150-D001, 40150-D003, and 40150-D006). The anti-polyhistidine antibody that was used in polyhistidine-mediated S1 protein immobilization (see Figure S1) was purchased from Thermo Fisher (MA1-21315). The horseradish peroxide (HRP) conjugated secondary antibody used in the IgG detection experiment was from the detection antibody in Thermo Fisher’s human total IgG ELISA kit (88-50550-22). The HRP conjugation of CR3022, D001, D003, and D006 antibodies were carried out with Abcam’s HRP conjugation kit (ab102890) with a molar ratio of antibody : HRP = 1 : 4.

### ELISA protocols

The illustrations for the reactor preparation protocol and the ELISA protocols can be found in Figure S2. For all steps, the working solution of the wash buffer and reagent diluent were diluted with UltraPure™ DNase/RNase-free distilled water to achieve 1× working concentration.

In the first step of the reactor preparation process (Figure S2(A)), the working solution of the capture antibody (*i.e*., D001 in S1 detection experiments) or the anti-His antibody (in IgG detection and antibody affinity experiments) were prepared by diluting the stock solution with the ELISA coating buffer (1× PBS, pH = 7.4) to achieve final concentrations of 10 μg/mL. Note that for the anti-S1 IgG detection and antibody affinity experiments, the second incubation step was used for blocking plus S1 protein immobilization (with 2 μg/mL of S1-His protein dissolved in 1% BSA). For the S1 protein detection, this step is used for blocking only (with 1% BSA in 1× PBS).

For the anti-S1 IgG detection experiments (see Figure S2(B) for the detailed protocol), various concentrations of monoclonal antibodies were prepared by diluting the stock solutions with 50 times diluted human serum (the serum was diluted with 1× reagent diluent, which correlates to 1% BSA). The working solution of the detection antibody (in this case, the HRP-conjugated secondary antibody) was prepared by diluting the stock solution 250 times in 1× reagent diluent.

For the S1 detection experiments (see Figure S2(C) for the detailed protocol), various concentrations of the S1 proteins were prepared by diluting the stock solution with 10 times diluted human serum. The working solution of the detection antibody (*i.e*., HRP-conjugated D003) was prepared by diluting the stock solution 1000 times in 1× reagent diluent. The final concentration of the detection antibody was 1 µg/mL.

For the antibody affinity experiments, various concentrations of HRP-conjugated monoclonal antibodies were prepared by diluting the stock solution in 1× reagent diluent.

### ELISA measurements

The signal intensities of the microfluidic ELISA were measured with the chemiluminescent imaging method, using a CMOS camera (see Figure 1(B) for an example of the chemiluminescent signals). To enhance the dynamic ranges of the ELISA, multiple exposures with adjustable exposure time were applied. All chemiluminescent intensities (CL intensity) were normalized to three seconds of exposure time. Detailed explanations about the chemiluminescent imaging and the multiple exposure approaches can be found in our previous publications^18^.

## Results

### Detection of anti-SARS-CoV-2 S1 IgG

The receptor binding domain (RBD) in the spike protein (located in the S1 subunit) on a coronavirus is responsible for binding to the membrane receptors on the target cell. It plays a critical role in the coronavirus cell entry process. For a patient infected by a coronavirus, his/her immune system will develop antibodies to bind and block the RBD of that specific coronavirus. Out of all types of neutralization antibodies, IgG has the longest lifetime in a person’s circulating blood. It is used as a biomarker for the evaluation of patient’s adaptive immune responses and the degree of recovery). In addition, as the major active ingredient, the concentration of SARS-CoV-2 S1 specific IgG can be used as the indicator for the strength of the convalescent plasma^19-23^.

For these reasons, IgG that binds specifically to SARS-CoV-2 S1 protein (especially the RBD) is the first biomarker that we aim to detect. The mechanism of IgG detection is illustrated in Figure 2(A). First, S1 protein is immobilized on the capillary inner surface through a poly-histidine mediation approach (see Figures S1 and S2(A) for details). Then, S1 specific IgG in the sample (such as serum) is attracted to the surface through immunosorbent reaction. Finally, the HRP-conjugated detection antibody is used to visualize the binding of the immobilized IgG. To ensure detection specificity, a monoclonal detection antibody that binds specifically to the Fc domain on human IgG was used. According to the protocol in Figure S2(B), the entire assay was completed in 15 minutes.

**Figure 2.**
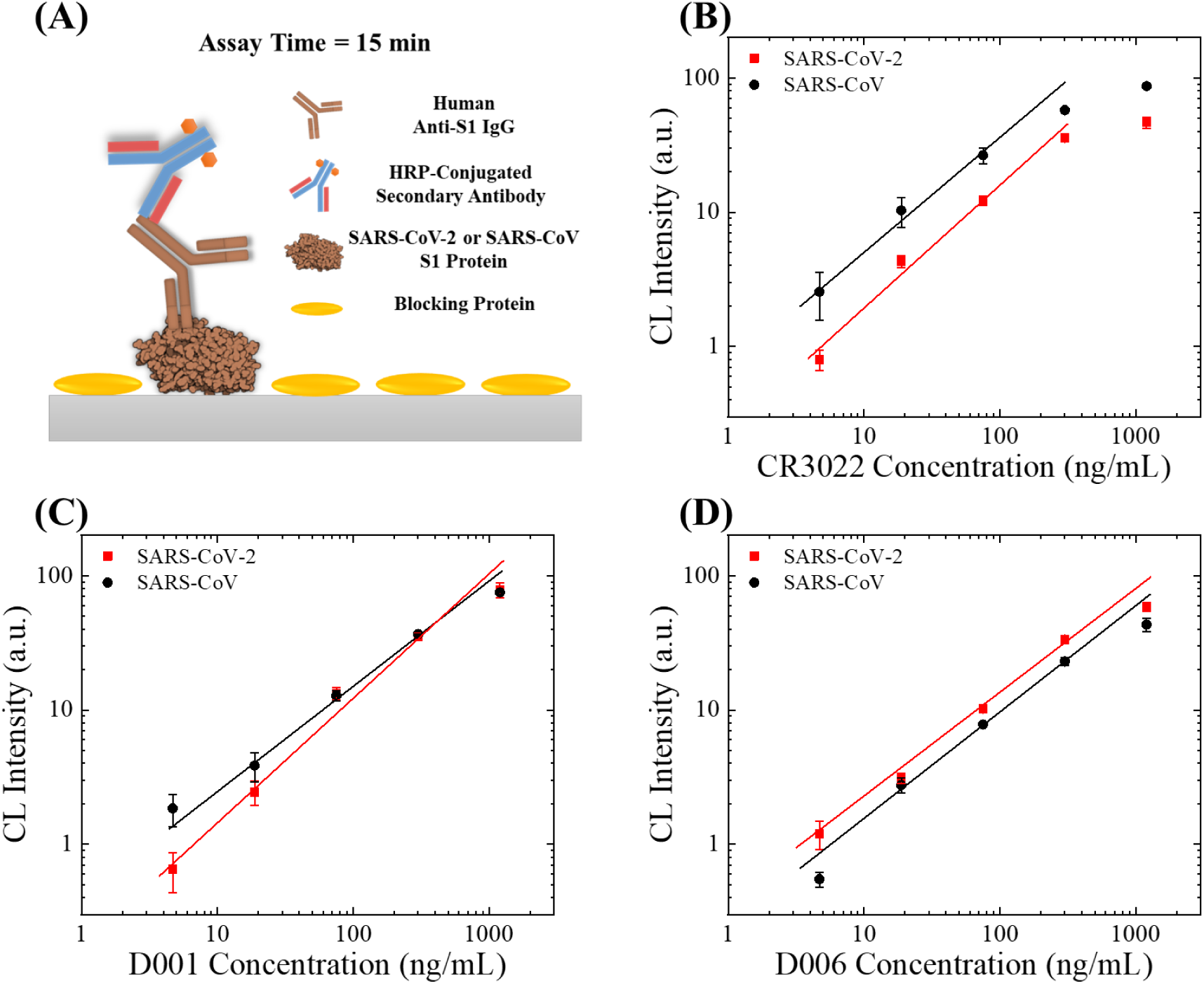
Detection of anti-S1 IgG. (A) Illustration of the assay mechanism. The sample-to-answer time of this assay is 15 minutes. (B)-(D) Detection of S1 specific IgG in 50 times diluted serum, against the S1 protein from SARS-CoV-2 (red squares) and SARS-CoV (black circles). The calibration curves are generated with three different monoclonal humanized antibodies (CR3022 in (B), D001 in (C), and D006 in (D)). The solid lines are the linear fit for the data in the log-log scale. Error bars are generated from duplicate measurements. See also Figure S3 for the entire dynamic range of CR3022, D001, and D006, and their respective lower limits of detection.

To validate the feasibility of our assay, we selected three recombinant and humanized monoclonal anti-SARS-CoV-2 S1 antibodies as the positive control antibodies. The first antibody, CR3022, is a therapeutic human IgG originally developed against the S1 protein of SARS-CoV. It was recently reported to have cross-reactivity towards the S1 protein of SARS-CoV-2^24-26^. The second and the third antibodies, D001 and D006, are humanized chimeric IgGs (the precursors of D001 and D006 were originally raised in mouse and rabbit, respectively) that are specific to the S1 protein of SARS-CoV. Based on the preliminary results conducted at Sino Biological internally, they were also believed to have high binding affinities towards the S1 protein of SARS-CoV-2. In order to mimic the actual clinical situations, we decided to use 50 times diluted human serum as the solvent of the IgG antibodies (50-1000 are the typical dilution factors of serum in actual serological analyses). To evaluate the differences in antibody’s affinity towards SARS-CoV-2 S1 and SARS-CoV S1, we performed a side-by-side study with these two types of S1 proteins for all three clones of antibodies.

The corresponding results are shown in Fig. 2(B)-(D). In general, the chemiluminescent intensities are proportional to the concentration of the spiked-in monoclonal antibodies. All these three antibodies are still detectable at 4.7 ng/mL with both types of S1 proteins. For D001 and D006, the signal for both types of S1 proteins is very similar, indicating that the antibodies’ binding affinity towards SARS-CoV-2 S1 and SARS-CoV S1 should be very similar (D006 may have a slightly higher affinity towards SARS-CoV-2 S1 than SARS-CoV S1). However, for CR3022, the signal for SARS-CoV-2 S1 is systematically lower than that for SARS-CoV S1, indicative of a weaker affinity of CR3022 towards SARS-CoV-2 S1 than SARS-CoV S1, which agrees with recently published findings about CR3022’s binding ability^24, 26^.

The entire dynamic ranges of these three antibodies against SARS-CoV-2 S1 can be found in Fig. S3. The linear dynamic range in the log-log scale for CR3022, D001, and D006 are 7-500 ng/mL, 2-1000 ng/mL, and 2-1000 ng/mL. The corresponding slope in the linear range is 0.64, 0.83, and 0.81, respectively. As a negative control, a human IgG isotype does not generate any detectable signal within the entire range of detection (0.7-4800 ng/mL). Due to the narrow linear dynamic range and relatively low binding affinity, CR3022 should not be used as the calibration standard of anti-SARS-CoV-2 S1 IgG. The remaining two antibodies, D001 and D006, are nearly the same in terms of the dynamic range and affinity towards SARS-CoV-2 S1. However, D006 has a better specificity for SARS-CoV-2 S1 compared to SARS-CoV S1. Therefore, D006 may be a better candidate as a calibration antibody.

### Detection of SARS-CoV-2 S1 protein

In addition to anti-S1 IgG, S1 protein itself may also be a marker in the prognosis of COVID-19. It may appear in blood for the patients who develop coronavirus viremia and the people who receive certain types of coronavirus vaccines (especially subunit vaccines). Recent evidence indicates that the viral proteins may also exist in mucus samples such as the saliva. To detect the S1 protein, we employed a standard sandwich ELISA format, as illustrated in Fig. 3(A), in which a monoclonal SARS-CoV-2/SARS-CoV S1 RBD-specific antibody (D001) was used as the capture antibody and another S1 RBD-specific antibody (D003) was used as the detection antibody SARS-CoV-2/SARS-CoV. To reduce the number of steps as well as the total assay time, we directly conjugated HRP molecules on the detection antibody. According to the protocol in Figure S2(C), the entire assay was completed in 20 minutes.

**Figure 3.**
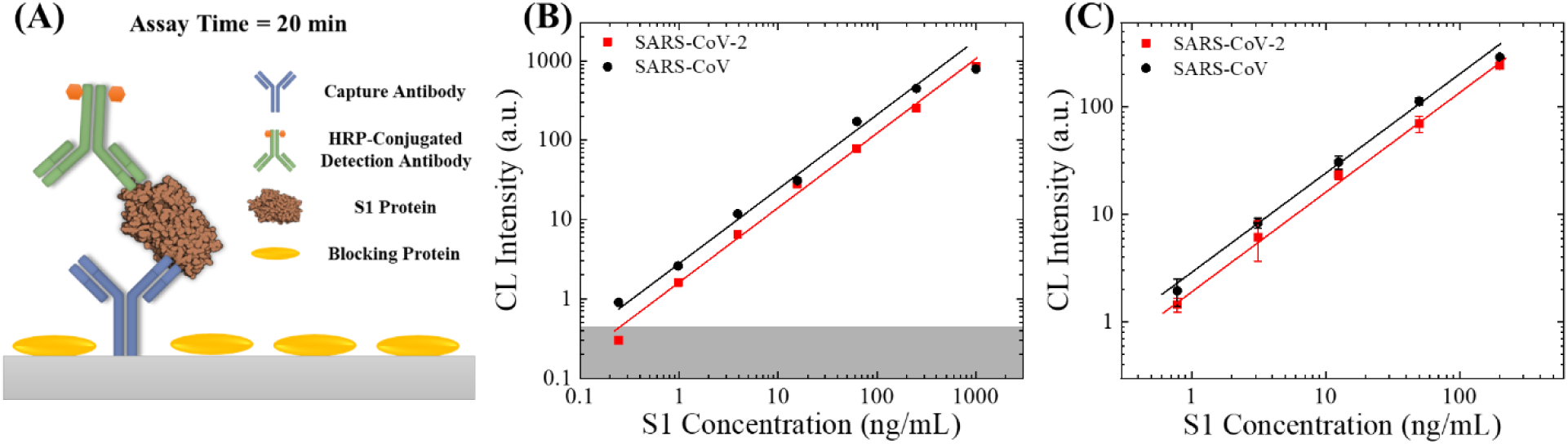
S1 protein detection. (A) Illustration of the assay mechanism. The sample-to-answer time of this assay is 20 minutes. (B) Entire dynamic ranges of SARS-CoV-2 S1 protein (red squares) and SARS-CoV S1 protein (black circles) in 10 times diluted human serum. The averaged background is subtracted from all data points. The solid lines are the linear fit of the data in the log-log scale. The grey shaded area marks 3×standard deviation of the background. The lower limit of detection (LLOD) for SARS-CoV-2 S1 protein and SARS-CoV S1 is 0.4 ng/mL and 0.2 ng/mL, respectively. (C) Calibration curves for S1 proteins between 0.78 and 200 ng/mL. The error bars are generated from duplicate measurements.

Same as in the IgG detection experiment, we performed a side-by-side study with the S1 proteins from SARS-CoV-2 and SARS-CoV. To mimic actual clinical settings, we used 10 times diluted human serum as the solvent of the S1 protein, as we do not expect to see a high concentration of viral S1 protein in serum (or saliva). The entire dynamic range of the S1 detection assay is presented in Fig. 3(B). The linear dynamic range for SARS-CoV-2 S1 and SARS-CoV S1 is 0.4-1000 ng/mL and 0.2-250 ng/mL with a slope of 0.89 and 0.92 (in the log-log scale), respectively. According to Figures 3(B) and (C), the detection of SARS-CoV S1 appears to have a higher sensitivity than SARS-CoV-2 S1. This is because both of antibodies (D001 and D003) used in this assay were originally raised against the RBD of SARS-CoV S1. A higher sensitivity in detecting SARS-CoV-2 S1 protein may be achieved in the future with the antibodies specifically developed against the S1 protein (or S1 RBD) of SARS-CoV-2.

#### Rapid affinity evaluation of monoclonal antibodies against SARS-CoV-2/SARS-CoV S1

Based on the studies in the previous sections, it is obvious that a good calibration standard (monoclonal human or humanized IgG towards SARS-CoV-2 S1 protein) is essential for performing quantitative evaluation of the patient’s immune response. High affinity antibodies are also essential for building a sensitive SARS-CoV-2 S1 ELISA kit. However, due to nascence of SARS-CoV-2 there is no “gold standard” antibody that can be used in IgG calibration or S1 antigen detection yet. Consequently, it is urgent to find humanized antibodies with high binding affinities. Unfortunately, traditional antibody evaluation approaches, such as plate-based ELISA, bio-layer interferometry (BLI), and surface plasmonic resonance (SPR), suffer from long assay time (2-3 hours for plate-based ELISA), small dynamic range (2 orders of magnitude for plate-based ELISA), and large sample concentration and consumption (10-100 µg/mL for BLI or SPR) ^27-29^. To address these problems, here we present a simple and rapid approach for the affinity assessment of monoclonal antibodies.

The assay mechanism and the corresponding protocol are illustrated in Figures 4(A) and S2(D), respectively. Our assay follows a single-step ELISA format. Same as in the IgG detection experiment, recombinant S1 proteins were first immobilized on the supporting surface (2 µg/mL), followed by the binding of IgG. To reduce potential sources of errors, we directly conjugated HRP molecule with purified IgG molecules (molar ratio IgG : HRP = 1 : 4, with the HRP conjugation kit from Abcam). The antibodies were then diluted to six different concentrations (1000 – 1 ng/mL with 4× serial dilution) and then applied to the S1 protein-coated ELISA reactors. The immobilized IgG can be quantified after a short 6 minutes of incubation and 4 times of rinsing. Although the antibodies may not reach equilibrium by the end of the incubation, the quantity of the immobilized antibody (also the signal intensity) should still have a positive correlation with the affinity of the antibody.

**Figure 4.**
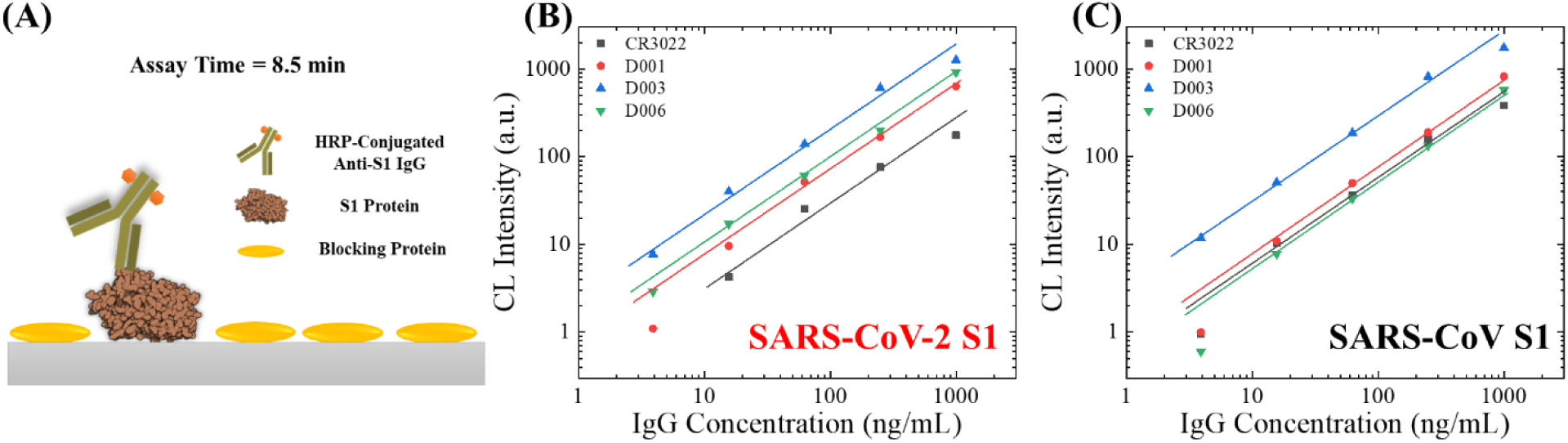
Antibody affinity screening. (A) Illustration of the assay mechanism, which uses a single-step ELISA. The sample-to-answer time of this assay is 8 minutes. (B)-(C) Calibration curves of 4 different monoclonal humanized S1 specific IgG against the S1 protein from SARS-CoV-2 (B) and SARS-CoV (C). The solid lines are the linear fit of the data in the log-log scale.

To compare the antibody’s affinity towards the S1 protein of SARS-CoV-2 and SARS-CoV, we performed a side-by-side experiment for all antibodies under test^30^. The antibodies were CR3022, D001, D006, and D003. The first three antibodies were used as calibration standards in the IgG detection experiments and D003 was used as the detection antibody in the S1 detection experiments. Thanks to the simple protocol, our assay exhibits excellent intra-assay consistencies (see Fig. S4 as an example), thus ensuring highly reliable measurements. As shown in Figure 4(B), these four antibodies have very different affinities towards the S1 protein of SARS-CoV-2. Note that the data for 1 ng/mL are not presented because the signal is not detectable for the three antibodies except D003 (the signal for D003 is very weak, see Figure S4). For the points within the linear dynamic ranges, the signal intensities with the strongest antibody (D003) are 6-10 times higher than the weakest antibody (CR3022). For example, the signal intensities for these two antibodies at 250 ng/mL are 612 and 75.9, respectively. On the other hand, the lowest detectable concentration for D003 is 1 ng/mL and the lowest detectable concentration for CR3022 is 16 ng/mL. This can be another evidence for the difference in antibody’s affinity. These observations agree with several recently published experiment results and our own preliminary results^24-26^ and our measurements at Sino Biological (the equilibrium dissociation constant, KD, for CR3022 and D003 is 6-12 nM and <1 nM, respectively, based on BLI measurements.

In contrast, as shown in Figure 4(C) the antibodies’ affinity towards SARS-CoV S1 can be very different from SARS-CoV-2 S1. For example, CR3022’s binding affinity towards SARS-CoV S1 is stronger than its affinity towards SARS-CoV-2 S1. Conversely, D006’s binding affinity towards SARS-CoV S1 is weaker than SARS-CoV-2 S1. In addition, the pattern of calibration curves with SARS-CoV-2 S1 is significantly different from that with SARS-CoV S1. While D001, D006, and CR3022’s affinities towards SARS-CoV-2 S1 vary significantly, they appear to be very similar to each other towards SARS-CoV S1. Although our current method allows us to rapidly evaluate the relatively affinity among all antibodies, it is unable to extract the exact value of KD. This problem can be resolved by introducing multiple calibration antibodies with known affinities.

## Discussion and Conclusion

We have demonstrated a portable chemiluminescent microfluidic ELISA system that is able to conduct sensitive detection and quantification of SARS-CoV-2-related biomarkers in only 15-20 minutes using micro-liter sized sample volumes. The LLOD of 2 ng/mL for IgG in serum was achieved using the humanized chimeric antibodies as the model system. We also successfully characterized different antibodies and identified an antibody candidate, D006, which can be used as the calibration antibody for quantitative evaluation of anti-SARS-CoV-2 S1 IgG. This approach can also be extended to evaluation of therapeutic convalescent plasma. Furthermore, we demonstrated sensitive detection of SARS-CoV-2 S1 protein (antigen) with the LLOD of 0.4 ng/mL. Finally, we showed that our technology can be used as an alternative approach for rapid (8.5 minutes) screening and validation of monoclonal anti-S1 antibodies. Our method requires only tens of nanograms, which is several orders of magnitude smaller than used traditional label-free methods (such as BLI and SPR), and will be useful in screening and selection of high affinity therapeutic neutralization antibodies and research-use antibodies^23, 27-29^.

We will continue to optimize our assays in multiple aspects. For IgG detection, we will investigate more humanized or patient-derived SARS-CoV-2 S1 antibodies to identify optimal calibrators with a high binding affinity and a large linear dynamic range. The differences in the antibodies’ affinity against monomeric S1 (most commonly used in antibody and vaccine development) and trimeric S1 (the natural conformation of S1 on SARS-CoV-2 virus) can also be explored^6^. For S1 protein detection, we will improve the detection sensitivity, since the abundance of SARS-CoV-2 S1 may be very low in actual patient samples. Based on our previous publications^17, 28^, the LLOD can be greatly enhanced when we replace the HRP-conjugated detection antibody with a biotinylated detection antibody once it become available. For antibody affinity evaluation, we aim to further optimize the assay’s protocol (such as adjusting the incubation time and rinsing time) and employ it to evaluate more therapeutic and research use antibodies.

Our approach also opens a door for other COVID-19 related clinical or laboratory researches. For example, the diagnostic value of the COVID-19 related biomarkers in serum and saliva (especially S1 specific IgA, IgM, and viral antigens such as the S1 and nucleocapsid (N) proteins) is currently under intensive evaluation^8, 31, 32^. The IgG detection method described in this work can be easily adapted to detect and quantify other types of antibodies such as IgM and IgA^31, 33, 34^. The concept for SARS-CoV-2 S1 protein detection can also be adapted to detect other types of viral antigens (such as the SARS-CoV-2 N protein), as described in Figure S5. Direct detection of viral antigens in patient samples such as serum and saliva may facilitate the rapid and cost-effective diagnosis of COVID-19^7, 35^. Finally, the microfluidic ELISA platform can be used to study the neutralization efficacy of therapeutic antibodies^23^ (see Figure S5), as well as for the recognition, evaluation, and phenotyping of natural and recombinant (fake) viral particles^36^.

## Acknowledgement

The authors thank the financial support from the Department of Biomedical Engineering.

