## Supplementary Information for "Rapid and quantitative detection of COVID-19 markers in micro-liter sized samples"

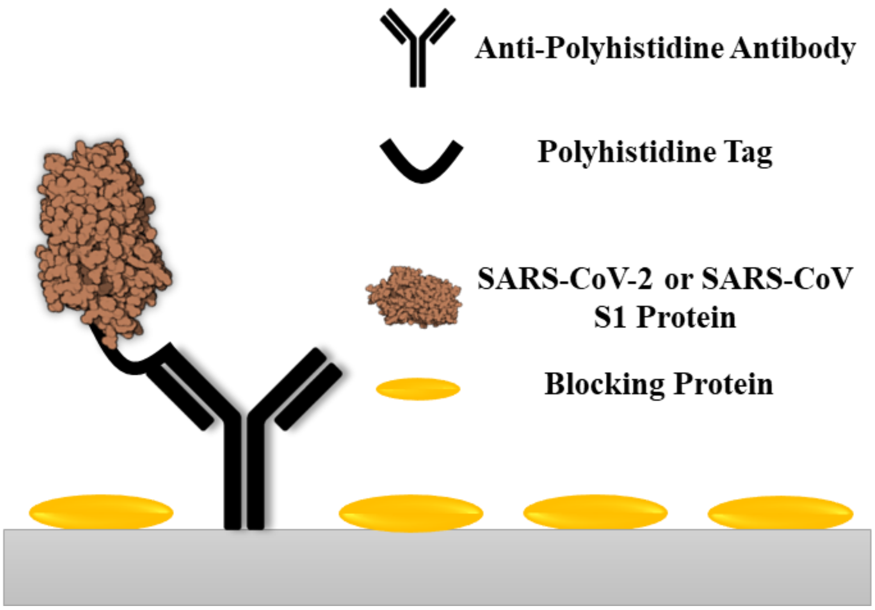


**Figure S1.** Conceptual illustration of polyhistidine-mediated S1 protein immobilization. The ELISA reactors were first coated with anti-polyhistidine antibodies. The polyhistidine tag (His-tag) on the recombinant S1 proteins can then bind to the immobilized anti-His tag antibodies. Since the His tag is located at the C- terminus of the S1 protein, the entire RBD should be accessible for antibody binding.

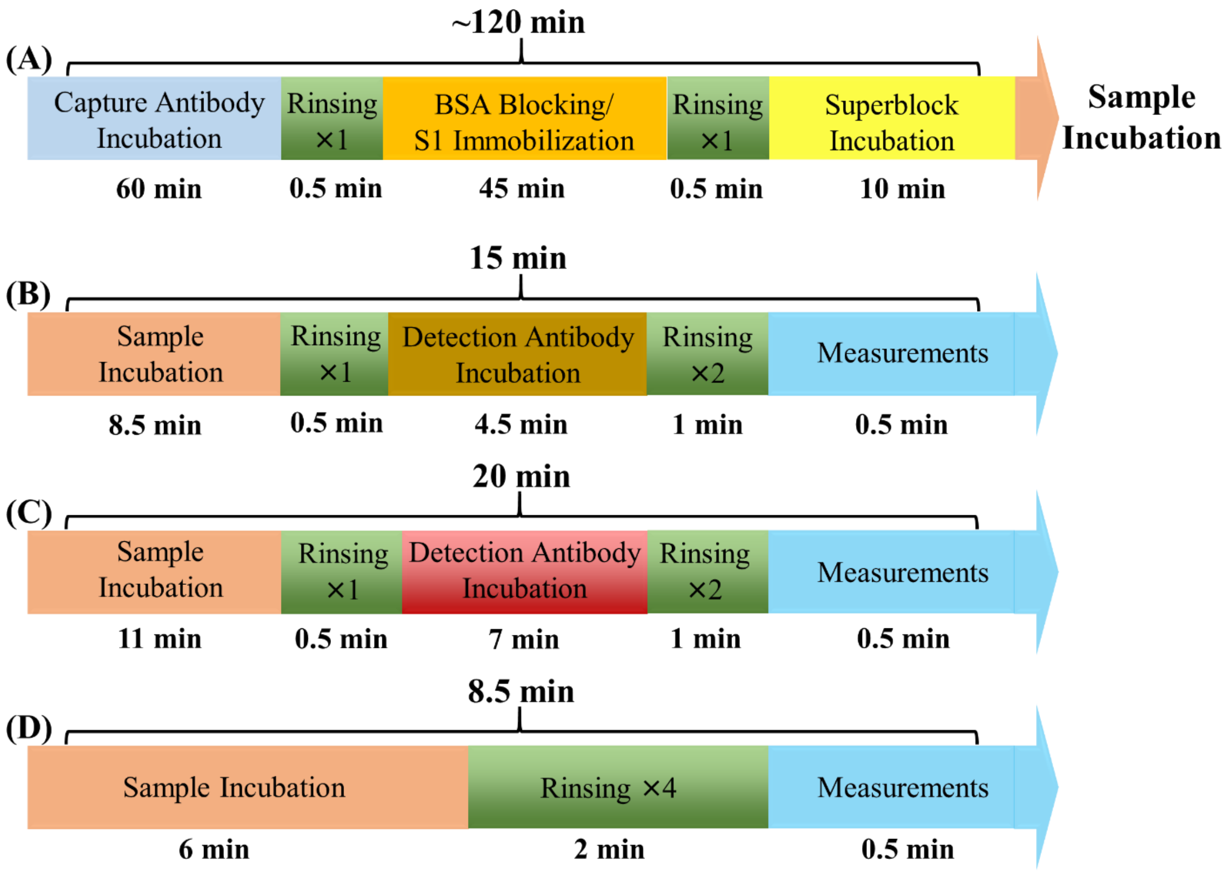


**Figure S2.** Graphical illustration for the assay protocols. (A) Procedure of the reactor preparation processes. For the anti-S1 IgG detection and antibody affinity evaluation experiments, the second incubation step is used for blocking plus S1 protein immobilization. For the S1 protein detection, this step is used for blocking only. The remaining steps are common for all three experiments. Note that the reactor preparation process can be done en-masse well in advance. (B) Protocol for the anti-S1 IgG detection. The assay time is 15 minutes. Note that the detection antibody is an anti-human IgG (Fc specific) secondary antibody. (C) Protocol for the S1 detection experiment. The assay time is 20 minutes. (D) Protocol for antibody affinity evaluation. The assay time is only 8.5 minutes.

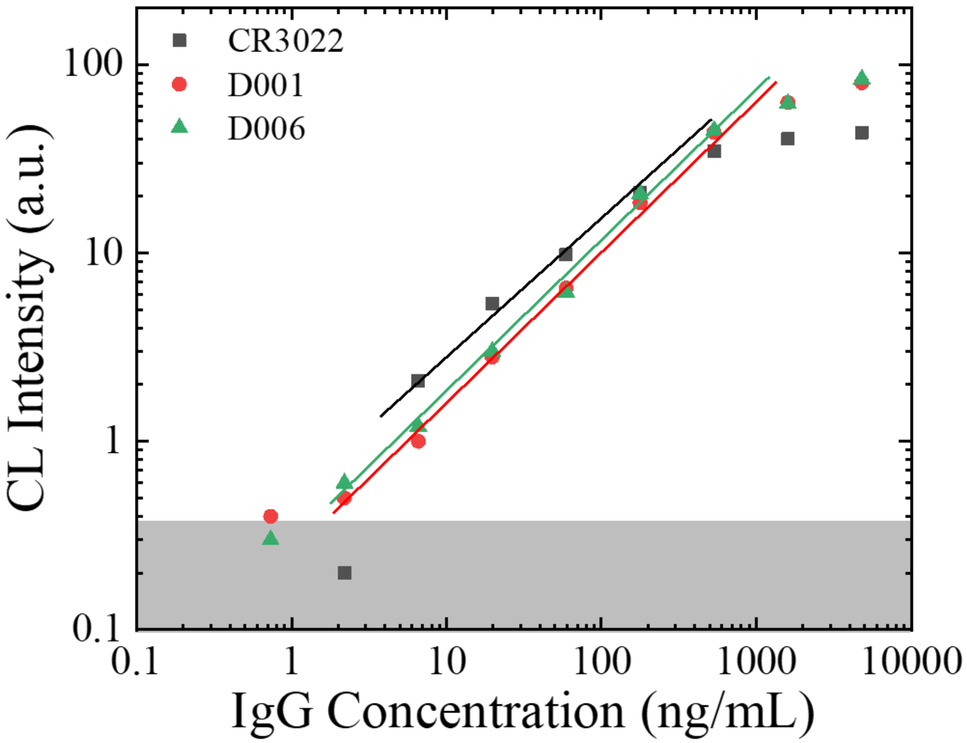


**Figure S3.** Entire dynamic ranges for the detection of the three humanized monoclonal antibodies. The concentrations were prepared from 3 times of serial dilution (starting from 4800 ng/mL). The averaged background is subtracted from all data points. The solid lines are the linear fit of the data in the log-log scale. The grey shaded area marks 3× standard deviation of the background. The LLODs for D001 and D006 are 2 ng/mL and the LLOD for CR3022 is 7 ng/mL. The linear dynamic range for D001, D006, and CR3022 is 2-1000 ng/mL, 2-1000 ng/mL, and 7-500 ng/mL, respectively. Due to the narrow linear dynamic range, CR3022 should not be used as the calibration standard of anti-SARS-CoV-2 S1 IgG.

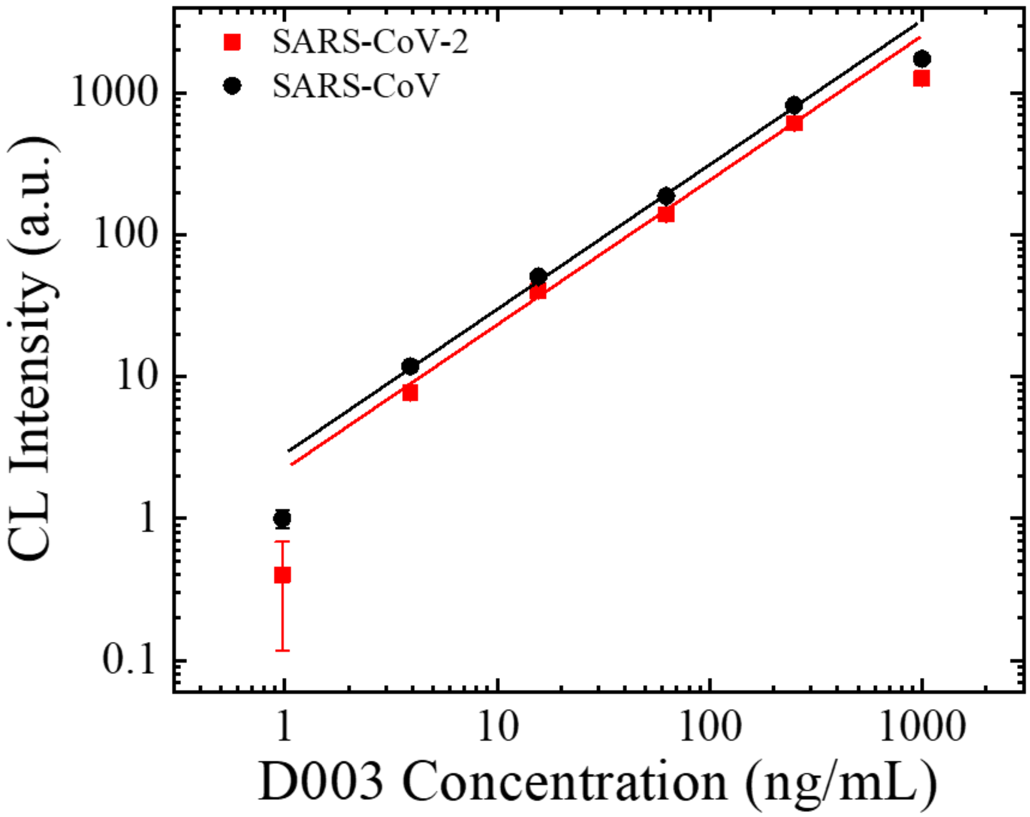


**Figure S4.** Dynamic ranges of a representative antibody, D003, in the antibody screening experiments. The data points fall out of the linear dynamic range at 1 ng/mL. The error bars are obtained from duplicate measurements. Due to the small intra-assay variances, the error bars are not visible in most datapoints.

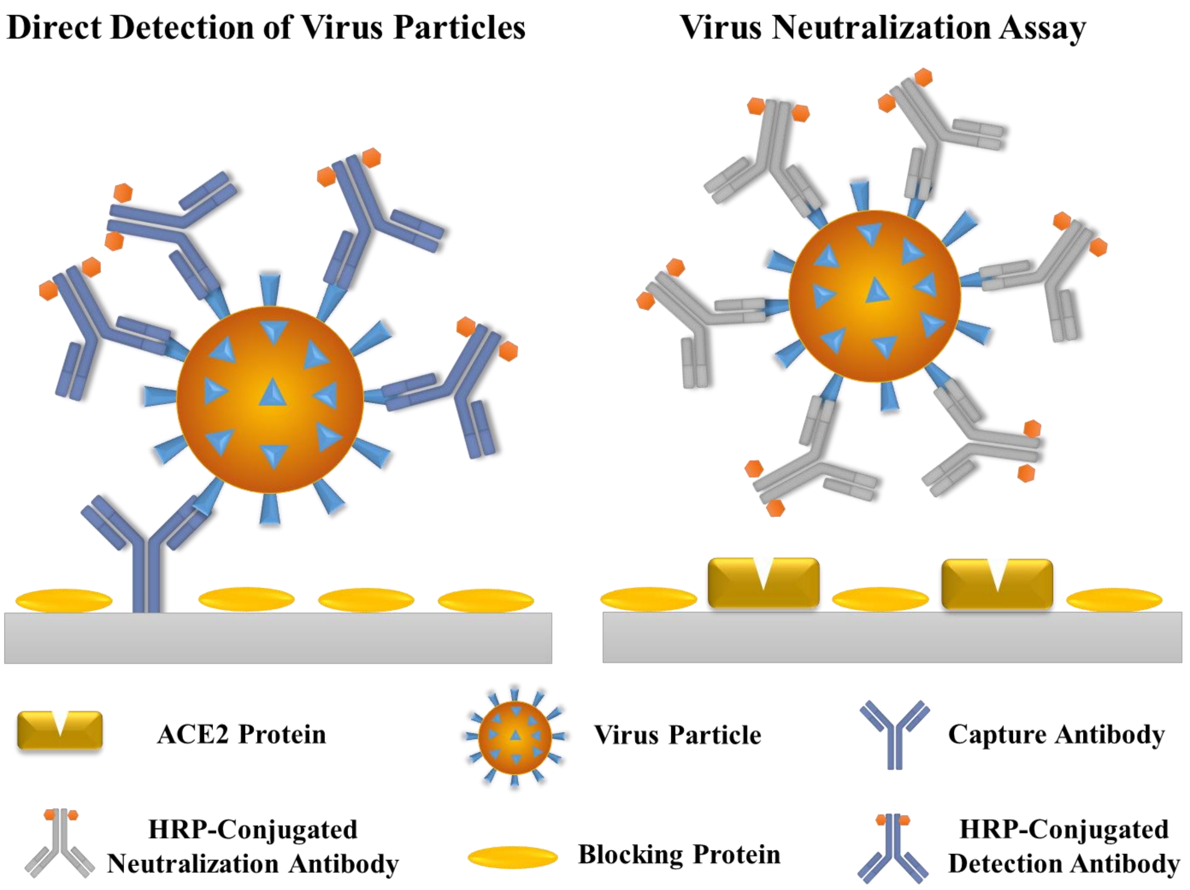


**Figure S5.** Potential COVID-19-related applications with our rapid microfluidic ELISA system, including direct detection of virus particles and virus neutralization assays (evaluation of neutralization antibodies). For direct detection of virus particles, since viruses are complex particles with multiple identical epitopes, the capture antibody and the detection antibody can be the same.
